# Visualization and Analysis of the Interaction Network of Proteins Associated with Blood-cell targeting Autoimmune Diseases

**DOI:** 10.1101/763672

**Authors:** Athina I. Amanatidou, Katerina C. Nastou, Ourania E. Tsitsilonis, Vassiliki A. Iconomidou

## Abstract

Blood-cell targeting Autoimmune Diseases (BLADs) are complex diseases that affect blood cell formation or prevent blood cell production. Since these clinical conditions are gathering growing attention, experimental approaches are being used to investigate the mechanisms behind their pathogenesis and to identify proteins associated with them. However, computational approaches have not been utilized extensively in the study of BLADs. This study aims to investigate the interaction network of proteins associated with BLADs (BLAD interactome) and to identify novel associations with other human proteins. The method followed in this study combines information regarding protein-protein interaction network properties and autoimmune disease terms. Proteins with high network scores and statistically significant autoimmune disease term enrichment were obtained and 14 of them were designated as candidate proteins associated with BLADs. Additionally, clustering analysis of the BLAD interactome was used and allowed the detection of 17 proteins that act as “connectors” of different BLADs. We expect our findings to further extend experimental efforts for the investigation of the pathogenesis and the relationships of BLADs.

## Introduction

Autoimmune diseases are a result of abnormal immune responses that fail to distinguish self from non-self. Despite their clinical diversity, they share one common characteristic, namely the excess activation of the immune system (1). The majority of them are multifactorial, since multiple genetic and environmental factors can trigger their onset. Historically, autoimmune diseases were considered to be rare, but epidemiological studies have shown that they affect 3-5% of the human population (2). It is remarkable that more than 100 distinct diseases (2), including multiple sclerosis and systemic lupus erythematosus, are classified as autoimmune diseases. However, an official classification of autoimmune diseases is still lacking.

Blood-cell targeting Autoimmune Diseases (BLADs), namely pernicious anemia (3), autoimmune neutropenia (4), autoimmune thrombocytopenic purpura (5), autoimmune lymphoproliferative syndrome (6), antiphospholipid syndrome (7), acquired aplastic anemia (8), paroxysmal nocturnal hemoglobinuria (9–11), primary acquired pure red cell aplasia (12,13) and autoimmune hemolytic anemia (14), are complex diseases that affect blood cell formation or inhibit blood cell production (more details about the description of each BLAD are given in Table 1). Interestingly, previous studies of *Conti et al.* (15), *Diz-Kucukkaya et al*. (16) and *Seif et al.* (17) have demonstrated clinical associations between some of the aforementioned BLADs. For example, autoimmune thrombocytopenic purpura (Evans syndrome) has been reported to coexist with paroxysmal nocturnal hemoglobinuria (15), antiphospholipid syndrome (16) and autoimmune lymphoproliferative syndrome (17).

**Table 1**: List of BLADs with the corresponding description, alternative names and PMIDs.

The genetic architecture that reflects the nature of autoimmune diseases results from the interaction of multiple genes and other factors. Genes associated with diseases encode proteins that interact within common functional modules (18). Proteins play a crucial role in molecular functions, hence, their interactions shape molecular and cellular mechanisms, which in turn control healthy and disease states in organisms (19). Nowadays, scientists are fascinated by protein-protein interaction (PPI) networks, since they provide useful, fast and inexpensive resources in the elucidation of disease mechanisms (20). The PPI networks are generally used for the identification of new candidate proteins associated with diseases (disease-related proteins hereafter) (18,21-25). As it has been proposed, one of the most effective ways to identify novel disease-related proteins is to find the interaction partners of known disease-associated proteins (21).

Even though their importance is well proven, computational approaches such as PPI networks have not been utilized extensively for the analysis of BLADs. To date, only a few studies have used a protein interaction network framework to obtain information regarding other autoimmune diseases, like Systemic Lupus Erythematosus (24), Rheumatoid Arthritis (26) and Celiac Disease (27). Moreover, discoveries of previous studies of *Conti et al.* (15), *Diz-Kucukkaya et al*. (16) and *Seif et al.* (17) motivated us to further study BLADs using a systemic approach. In this paper, a PPI network analysis was conducted to suggest candidate BLAD-related proteins and potentially identify essential proteins associating these diseases. Our approach combines information from PPI network analysis based on graph theory and autoimmune disease terms for the identification of candidate BLAD-related proteins. Furthermore, clustering analysis of the PPI network was used to uncover significant proteins of BLADs’ relationships.

## Methods

The research methodology used in this study involves the steps described below. Fig. 1 provides an overview of the methodology.

**Fig. 1:**
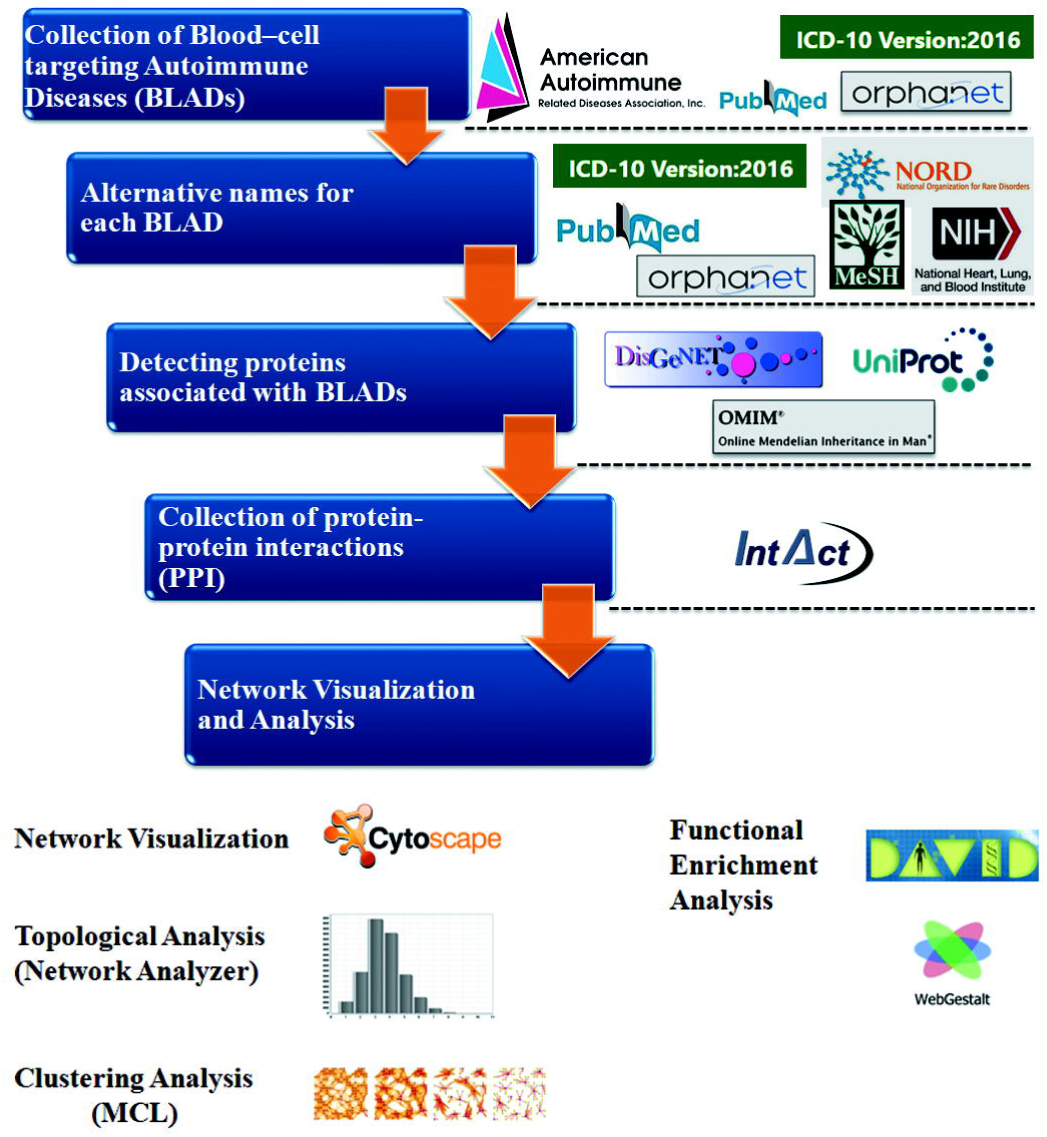
Workflow diagram.

### Collection of BLADs and their alternative names

Despite our efforts, we were unable to find an official classification of autoimmune diseases and it was thus necessary to exploit the existing list of Autoimmune Diseases by the American Autoimmune Related Diseases Association (AARDA) (https://www.aarda.org/). BLADs were selected on the basis of autoimmune responses against self-antigens of blood cells or their ability to hinder normal blood cell production.

The ICD-10 (28) and Orphanet (29) databases were used to find codes for autoimmune diseases, in order to ascertain that they are distinct diseases. At the same time a literature search was conducted to verify that these diseases belong to autoimmune diseases and which of them meet the above criteria.

However, it is becoming increasingly evident that disease nomenclature is changing over time. Due to this, a search for retrieving the official nomenclature of each BLAD from the following databases: ICD-10, Orphanet, NORD (30), MeSH (31) and NIH–NHLBI (32) was imposed. Subsequently, an extra step was taken for the verification of the attributed terms, through literature search.

### Detection of proteins associated with BLADs and their protein-protein interactions (PPI)

All selected alternative names were used to detect proteins associated with BLADs. These proteins can either be genetically associated with the diseases or present in high/low levels in patients’ plasma. For this purpose, disease-related protein association data were extracted from the OMIM (33) and DisGeNET (34) databases, which contain, among others, data of genes and variants associated to human diseases. A script was written in Perl to detect proteins associated with BLADs automatically in the UniProt human proteome (release date: 15-03-2017). Finally, all disease-related proteins gathered with the above method were manually verified for their association with BLADs.

Subsequently, using the UniProt Accession Numbers (ACs) of the disease-related proteins, a query in IntAct (35) – a database of highly curated experimental molecular interactions – was performed, in order to retrieve all known experimentally verified interactions of these proteins. All interactions between associated proteins and their first neighbors were collected. Interaction data were downloaded in a MI-TAB 2.7 format file (36). From this file all non-human interactions were excluded, as well as interactions with chemical compounds.

### Visualization and analysis of the PPI network

The PPI network was visualized and analyzed using the Cytoscape 3.4.0 (37), a well-known open source software platform for the visualization of molecular interaction networks (38). Subsequently, a **topological analysis** was performed with the evaluation of simple and complex parameters using the NetworkAnalyzer (39), a user-friendly inbuilt Cytoscape plugin.

Afterwards, **functional enrichment analysis** was performed with the use of two bioinformatics tools, DAVID (40) and WebGestalt (41). DAVID was used to detect statistically significant overrepresented Gene Ontology (GO) (42) terms in the network, and associate the network’s proteins with disease terms from Gene Association Database (GAD) (43) and KEGG pathways from KEGG PATHWAY Database (44). WebGestalt was used to detect the network proteins’ phenotype from Human Phenotype Ontology (HPO) (45). A significance was set at a P-value<0.05 for both aforementioned annotation tools (DAVID, WebGestalt).

Thereafter, the top scoring proteins (approximately the top 10% of the network’s proteins) were selected for each network centrality: degree, betweenness and closeness. An extra step for verifying the top scoring proteins included the collection of significantly enriched GO terms for: (a) the entire BLAD interactome, (b) high scoring proteins for each centrality measure and (c) proteins associated with BLADs. Then, disease ontology terms were associated with BLAD interactome’s proteins – where only proteins with highly statistically significant autoimmune disease ontology terms were collected. Subsequently, a Venn diagram was created in order to represent proteins with high scoring centralities, proteins associated with BLADs and proteins with significant autoimmune disease ontology terms. The **candidate BLAD-related proteins** were those that were present in both the lists of proteins with high scoring centralities and those with significant autoimmune disease ontology terms but did not belong to the list of proteins associated with BLADs.

Thereupon, a **clustering analysis** was performed with the utilization of the Markov Clustering algorithm (MCL) (46), a proper algorithm for finding densely connected regions in a network. Several studies have proved that MCL effectively identifies high-quality functional modules (47,48), and produces more accurate results for biological networks (49,50). The inflation value of MCL was set to 1.8, in compliance with other analyses that proved this value as suitable for biological networks (47,51,52).

Subsequently, our analysis was focused on clusters with at least 2 proteins associated with BLADs – in order to assess whether they interact with each other directly or through other proteins. In an effort to depict the type of connection between those proteins, the following orders of connection were used:

I. **1^st^ order connection:** proteins with direct interaction
II. **2^nd^ order connection:** proteins that have at least one common 1^st^ neighbor
III. **3^rd^ order connection:** proteins whose 1^st^ neighbors interact with each other
IV. **4^th^ order connection:** proteins that have at least one common 2^nd^ neighbor

The proteins which act as “bridges” in the connection of those associated with BLADs, will be henceforth termed “bridge proteins”.

## Results

### BLADs and their alternative names

The final list of BLADs is composed of 9 autoimmune diseases as presented in Table 1. ICD-10 codes were found for all diseases, except for autoimmune neutropenia, while Orphanet codes were retrieved for all of them (**Supplementary Table 1**). **Supplementary Figure 1** outlines an overview of the basic protocol used to collect BLADs.

Alternative names were retrieved for all autoimmune diseases, with the exception of acquired aplastic anemia (Table 1).

**Table 1**: List of Blood-cell targeting Autoimmune Diseases (BLADs) with the corresponding target cells, description/symptoms, alternative names and PMIDs.

### Construction and analysis of BLAD interactome

The initial data set of selected proteins associated with BLADs consists of 51 proteins (Table 2), 42 of which have PPI data deposited in IntAct (**Supplementary Table 2**). The PPI data were manipulated with the use of Cytoscape 3.4.0 to construct the PPI network, which is composed of 427 protein nodes and 1439 interaction edges (Fig 2). Cytoscape.js (53) was used to create interactive networks, available at http://83.212.109.111/BLAD. Detailed description of the functionalities offered by the interactive BLAD Interactome web application is available at **Supplementary File 1**. The PPI data obtained from IntAct are given in **Supplementary Table 3**.

**Table 2**: The data set of proteins associated with BLADs.

**Fig. 2:**
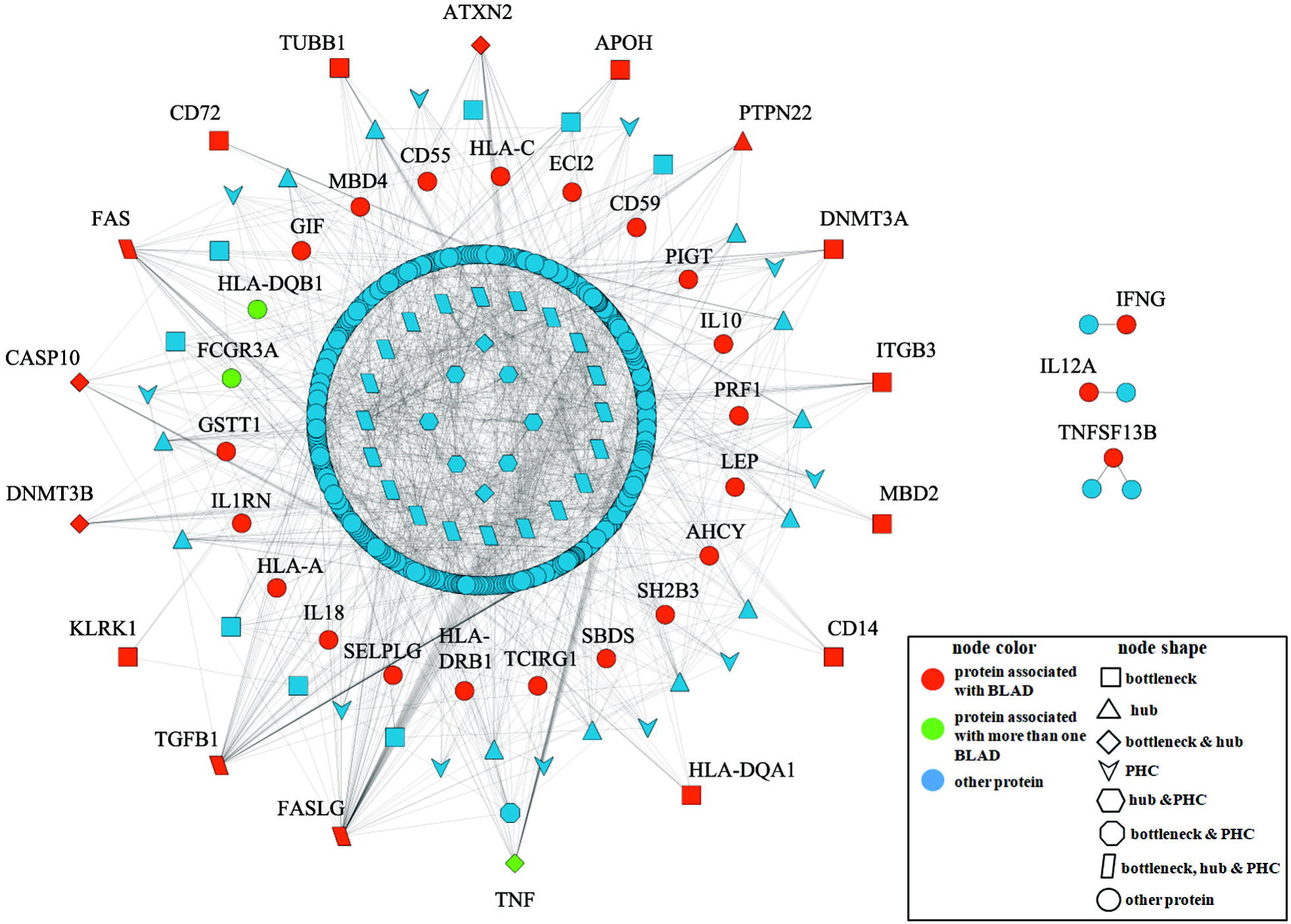
The BLAD interactome. This PPI network consists of 427 nodes (proteins) and 1439 edges (interactions). Red-colored nodes represent proteins associated with one BLAD, whereas green-colored nodes represent proteins associated with more than one BLAD. Blue-colored nodes are other proteins of the network. Bottlenecks and hubs are represented, respectively, as squares (■) and triangles (▲). Proteins which are both bottlenecks and hubs are shown as diamonds (♦). PHCs proteins are depicted as downward arrowheads 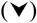. Proteins which are either hubs and PHCs or bottlenecks and PHCs are represented, respectively, as hexagons 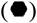 and octagons 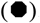. Proteins which are bottlenecks, hubs and PHCs are shown as parallelograms (?). For details regarding the proteins presented in this figure please refer to Supplementary Table 4. Interactive network available at http://83.212.109.111/BLAD.

Topological analysis is an efficient way to gain insight into a network’s topology and its participating proteins. After using NetworkAnalyzer, the network density, a metric that shows how sparse or dense a network is, has a value of 0.016. Also, the network’s tendency to be divided into clusters, represented by the clustering coefficient (CC) has a value of 0.142 and the characteristic path length (CPL) (54) is 3.532. The node degree distribution *P(k)* (55), follows the power-law P(k) − *Ak ^γ^*, where A is constant and γ is the degree exponent. In our case, the distribution is of the following form:

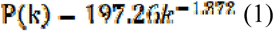

It is immensely significant for our analysis to decipher the critical players of our network, in accordance with the most important network centralities, namely degree, betweenness and closeness. *Hub* proteins can be identified by their high degree centrality, while *bottlenecks* are distinguished by high betweenness centrality (56). An indication of a fast information flow through a protein in a network, is its high closeness centrality. Proteins with that characteristic are hereby referred to as *PHCs (proteins with high closeness centrality)*. The proteins of the BLAD interactome (excluding the three isolated connected components) were sorted based on centrality measures and approximately the top 10% of the network’s proteins with the highest values were selected. Taking into account the overlapping proteins among the three protein lists, a total of 77 proteins were finally selected (**Supplementary Table 4**). Specifically, in the BLAD interactome, 13 proteins are *hubs* (Fig. 2, triangles), 17 proteins are *bottlenecks* (Fig. 2, rectangles), 6 proteins are *hubs* and *bottlenecks* (Fig. 2, diamonds), 11 proteins are *PHCs* (Fig. 2, V-shaped nodes), 6 proteins are *hubs* and *PHCs* (Fig. 2, hexagons), 1 protein is a *bottleneck* and *PHC* (Fig. 2, octagon), and 23 proteins are *hubs*, *bottlenecks* and *PHCs* (Fig. 2, parallelograms). Remarkably, 17 out of the 42 proteins associated with BLADs, present in our initial data set, have a significant role in the BLAD interactome.

### Functional enrichment analysis

To gain further insight into the function of the BLAD interactome proteins and the 42 proteins associated with BLADs, significantly related GO terms were found utilizing DAVID (**Supplementary Table 5**). Additionally, in order to verify the chosen network centralities’ ability to pinpoint significant related GO terms, functional enrichment analysis was performed for the top scoring proteins of each centrality measure. The most shared and statistically significant GO terms among these proteins are reported in **Supplementary Table 5**. The majority of proteins are implicated in several immune processes, such as T-cell co-stimulation (GO:0031295), extrinsic apoptotic signaling pathway (GO:0097191) and antigen processing and presentation (GO:0019882).

### Identification of candidate BLAD-related proteins

In order to find proteins among the 77 proteins with high scoring centralities – already associated with autoimmune diseases – we combined three protein lists: 77 proteins with high scoring centralities, 42 proteins associated with BLADs, and 120 proteins associated with significant autoimmune disease ontology terms. Through this combination, the list of proteins with high centrality scores emerged, which contained 77 proteins in total, out of which 17 proteins were associated with BLADs. Also, out of the 120 proteins associated with statistically significant autoimmune disease ontology terms, 32 proteins are associated with BLADs. It is worth mentioning that 14 proteins were found to be significantly associated with autoimmune disease ontology terms and have high centralities scores, as shown in the Venn diagram of Fig. 3 **(highlighted in yellow color)**. These proteins are depicted as black nodes in the BLAD interactome (Fig. 4). Additionally, all details regarding proteins belonging to at least 2 lists are given in **Supplementary Table 6**. Thus, it was feasible to identify 14 candidate BLAD-related proteins. The candidate BLAD-related proteins, the statistically significant autoimmune disease ontology terms, the interactions with proteins associated with BLADs, the three centrality measures and the corresponding ranks of the 14 important proteins are all shown in Table 3. Moreover, the annotation regarding 14 candidate BLAD-related proteins was based on EntrezGene database (57) focusing – when possible – to their participation in immune system activities (details in Table 3).

**Fig. 3:**
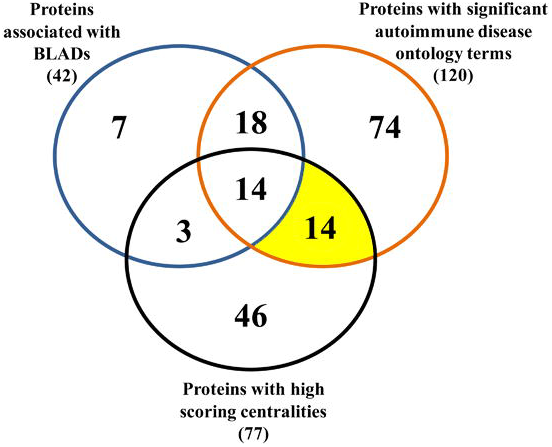
Venn diagram of candidate BLAD-related proteins. This diagram was created, using information about the proteins with high scoring centralities, significant autoimmune disease ontology terms and known proteins associated with BLADs. The yellow color highlights the 14 candidate BLAD-related proteins that have high protein network centrality scores and significant autoimmune disease ontology terms associations, and also do not belong to the list of already known BLAD-associated proteins. For details regarding the proteins presented in this figure please refer to Supplementary Table 6.

**Fig. 4:**
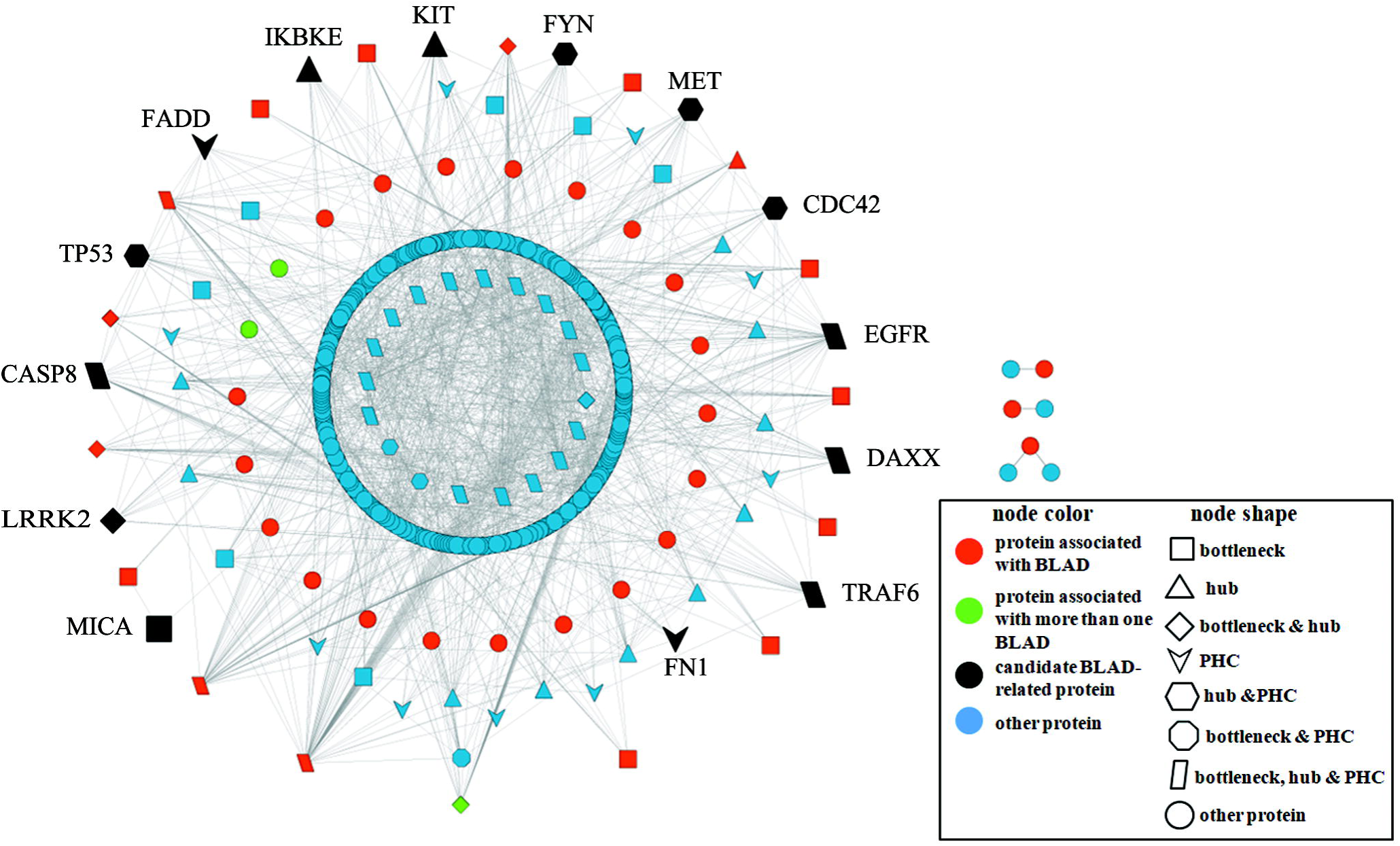
Visualization of 14 candidate BLAD-related proteins in the BLAD interactome. Red-colored nodes represent proteins associated with one BLAD, whereas green-colored nodes represent proteins associated with more than one BLAD. Black-colored nodes are the 14 candidate BLAD-related proteins, identified through our analysis.

**Table 3: Identification of candidate BLAD-related proteins**. The column “**Centrality measures”** presents proteins which are ranked in various centrality measures in the order: *Degree-D, Betweenness-B* and *Closeness-C*. The rank of each protein is given inside the braces of the corresponding centrality measure in the top 48 rankings (approximately the top 10% of the network’s proteins). Proteins ranking high in all three measures are shown first, followed by proteins with two measures and one measure, respectively. The column **“Autoimmune disease ontology terms”** represents the statistically significant autoimmune disease ontology terms per protein. The column “**Interactions with proteins associated with BLADs”** shows the interaction(s) of each protein with protein(s) associated with BLADs-whereas the corresponding association with BLAD can be seen inside the braces.

### Clustering and functional enrichment analysis

#### Clustering analysis

An efficient way to reduce the BLAD interactome’s complexity is the extraction of functional modules. For this purpose a clustering analysis was performed, utilizing MCL. The identification of functional modules is similar to finding communities (clusters) in a network (58). The BLAD interactome was divided in 42 clusters, 32 of which were composed of three or more proteins (Fig. 5), while 10 were not used for further analysis, since they contained only two proteins. The interconnections between proteins in three clusters consisting of at least two proteins associated with BLADs were the focal point of our subsequent analysis. Even though we chose to focus our analysis on these clusters, researchers are welcomed to extrapolate from the connections between proteins in the entire network or in specific clusters, based on their diseases or proteins of interest, by using our online web application available at http://83.212.109.111/BLAD/ (**Supplementary File 1**).

**Fig. 5:**
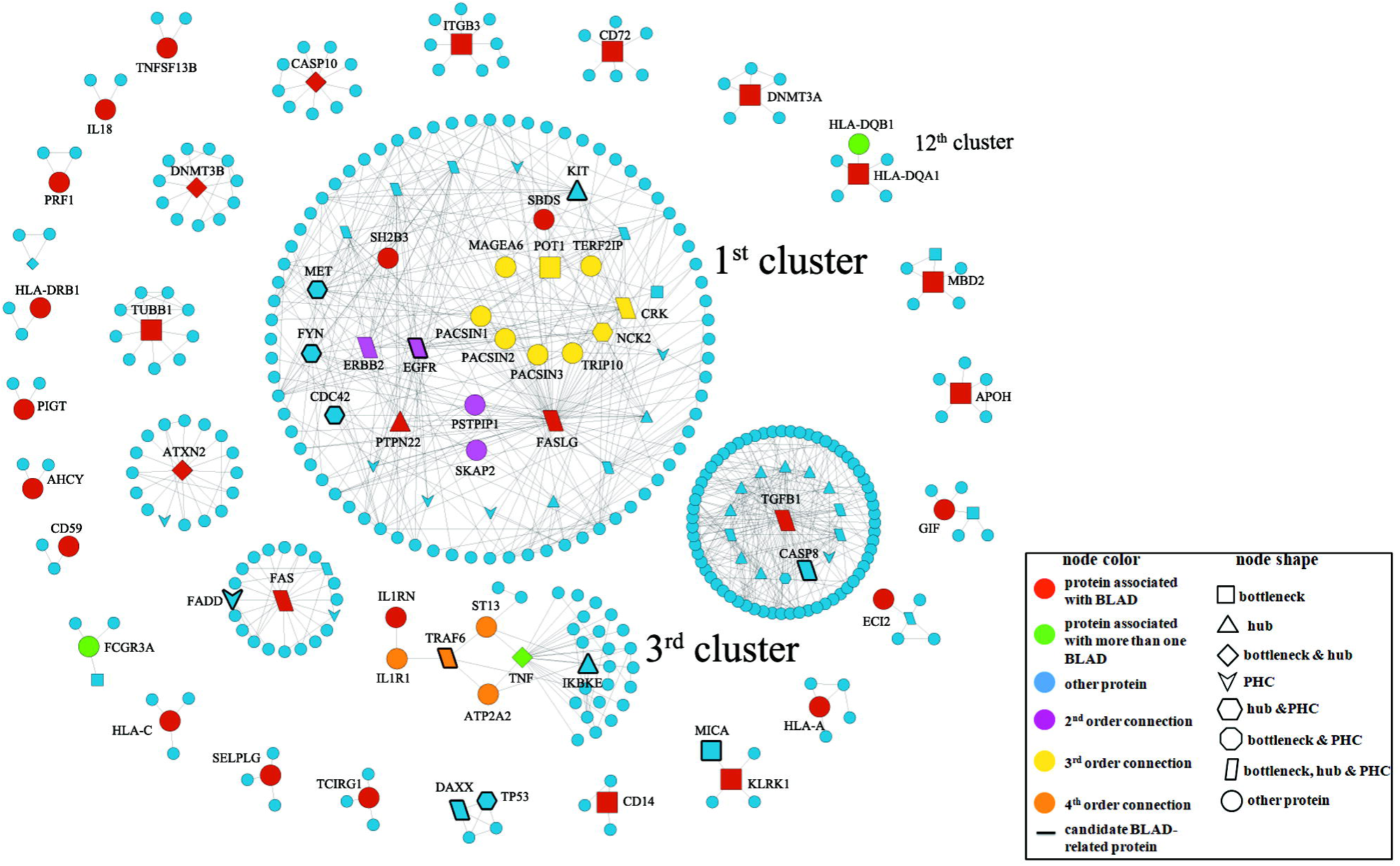
Clustering analysis of the BLAD interactome. In the first cluster, the 2^nd^ and 3^rd^ order connections among proteins associated with BLADs (red-colored nodes) are represented, respectively, as magenta- and yellow-colored nodes. In the third cluster, the 4^th^ order connection among proteins associated with BLADs (red-colored nodes) is depicted as orange-colored nodes. The thicker node outline shows the candidate BLAD-related proteins. Shapes and colors of all nodes are explained in the visual legend (bottom right). For details regarding the connections of proteins associated with BLADs presented in this figure, please refer to Supplementary Figures 2-5.

The first cluster consists of 106 proteins, including 4 proteins, SH2B3, PTPN22, FASLG, and SBDS (Fig. 5, **cluster 1-red nodes**), associated with antiphospholipid syndrome, autoimmune thrombocytopenic purpura, autoimmune lymphoproliferative syndrome, and acquired aplastic anemia, respectively. Notably, there are 13 proteins which act as “bridges” (Fig. 5, **cluster 1-magenta and yellow nodes**) among the aforementioned 4 proteins. Specifically, each pair of SH2B3-PTPN22 and PTPN22-FASLG proteins has a 2^nd^ order connection with 2 bridge proteins correspondingly (Fig. 5, **cluster 1- magenta nodes**). Furthermore, FASLG has a 3^rd^ order connection with SBDS protein, via 9 bridge proteins (Fig. 5, **cluster 1-yellow nodes**).

Subsequently, the third cluster integrates 32 proteins, including 2 proteins, TNF and IL1RN (Fig. 5, **cluster 3-green and red nodes**), the first associated with both autoimmune thrombocytopenic purpura and paroxysmal nocturnal hemoglobinuria, and the second with autoimmune thrombocytopenic purpura. Interestingly, it was observed that they have a 4^th^ order connection and there are 4 bridge proteins (Fig. 5, **cluster 3-orange nodes**) among them.

Finally, the twelfth cluster consists of 6 proteins, including 2 proteins – HLA-DQA1 associated with paroxysmal nocturnal hemoglobinuria, and HLA-DQB1 associated with autoimmune neutropenia and paroxysmal nocturnal hemoglobinuria – which have a 1^st^ order connection. Further details about proteins and their connections are reported in **Supplementary Figures 2-5**.

It is of utmost importance for our analysis to note, that the first cluster includes 5 proteins (Fig. 5, **cluster 1-nodes with thick black border)** out of the 14 candidate BLAD-related proteins, among them the bridge protein EGFR (Fig. 5, **cluster 1-magenta node with thick black border**). Additionally, in the third cluster 2 proteins are observed (Fig. 5, **cluster 3-nodes with thick black border)**, which belong to the data set of 14 candidate BLAD-related proteins, including the bridge protein TRAF6 (Fig. 5, **cluster 3-orange node with thick black border**). These proteins have high centrality scores and statistically significant associations with autoimmune disease ontology terms.

#### Functional enrichment analysis

For each cluster statistically significant GO terms were obtained, namely biological process, molecular function and cellular components. Concerning the clusters of our interest (1^st^, 3^rd^ and 12^th^), more details about biological processes, molecular functions and cellular components are shown in **Supplementary Table 7**, **Supplementary Table 8** and **Supplementary Table 9**, respectively.

#### Pathway analysis and disease association

The KEGG pathway analysis enlightens our knowledge about the common pathways in which the proteins of each cluster participate. In the 1^st^ cluster, the most highly statistically significant terms were detected. Those were focal adhesion (hsa04510), ErbB signaling pathway (hsa04012), T-cell receptor signaling pathway (hsa04660), and proteoglycans in cancer (hsa05205). Furthermore, in the 3^rd^ cluster the NF-kappa B signaling pathway (hsa04064) was dominant.

An extra disease association analysis was conducted - where significant associated diseases were identified for each cluster. Surprisingly, HIV and cutaneous squamous cell carcinoma are statistically significant for the 1^st^ cluster. Rheumatoid arthritis, sepsis and heart allograft rejection are highly related to proteins of the 3^rd^ cluster. The KEGG pathways and associated diseases for the 1^st^ and 3^rd^ clusters are given in **Supplementary Table 10** and **Supplementary Table 11**, respectively.

#### Human Phenotype Ontology (HPO) analysis

The HPO analysis was conducted with the goal of finding statistically significant phenotypic abnormalities of proteins in the three clusters of interest (1^st^, 3^rd^ and 12^th^). Abnormality of cellular immune system, skin physiology and myeloid leukocytes are highly related to the proteins of the 1^st^ cluster. Interestingly, common phenotypic abnormalities were detected for FASLG, PTPN22 and PSTPIP1 (**Supplementary Figure 2**), and these are abnormalities of immune system physiology and cellular immune system, increased inflammatory response, abnormality of blood and blood-forming tissues, and generalized abnormality of the skin. Also, abnormalities of the cellular immune system, skin physiology, leukocytes, blood and blood-forming tissues were found in common for PTPN22, SH2B3 and EGFR (**Supplementary Figure 3**). Additionally, abnormalities of immune system physiology, cellular immune system, blood and blood-forming tissues were identified for FASLG, SBDS and POT1 (**Supplementary Figure 4**). No statistically significant HPO terms were detected for other bridge proteins connecting FASLG and SBDS, except for POT1.

Regarding the 12^th^ cluster, the abnormalities of proteins characterize rheumatoid arthritis, keratoconjunctivitis sicca and achalasia. For the 3^rd^ cluster no statistically significant HPO terms were detected (**Supplementary Figure 5**). Further details regarding the 1^st^ and 12^th^ clusters are available in **Supplementary Table 12**.

## Discussion

In recent years, PPI networks have been used in targeting genes responsible for causing diseases and identifying new candidate disease-related proteins, in various disorders, among which are type II diabetes, primary immunodeficiencies, systemic lupus erythematosus, and other severe disorders (18,21-25). *In silico* methods have turned into a fast and inexpensive alternative way for the detection of proteins that could be considered as promising candidate drug targets (59). However, studies using PPI network analysis in relation to BLADs have not been performed to date. For this purpose, we performed an *in silico* analysis of the interaction network of proteins associated with BLADs, in order to identify candidate BLAD-related proteins, as well as proteins that may play an essential role in the relationships between these diseases.

From graph theory-based analysis, the topology of the network was extracted. Our network has a density of 0.016, which shows that the BLAD interactome is a sparsely connected network, in accordance with other biological networks (60). The clustering coefficient (CC=0.142) and the characteristic path length (CPL=3.532) are higher and lower, respectively, in comparison with those of a random network with the same number of nodes (CC=0.0023, CPL=7.12). The two criteria (high CC and less than expected CPL) reveal that the BLAD interactome has small-world properties (61), which means that proteins communicate quickly with each other, as exhibited by other PPI networks (62).

According to several studies, PPI networks are scale-free (63). The main characteristic of scale-free networks is that they follow the power law node degree distribution *P(k) (*P(k) = *Ak*^*−γ*^). The fact that our network also follows the power law distribution means that the majority of proteins in the BLAD interactome has few interactions with other proteins, while there exists a small number of proteins that have high degree connectivity called hubs (55). In an effort to rank the network’s proteins according to their importance, the degree, betweenness and closeness centralities were calculated for all BLAD interactome’s proteins.

The value of each network centrality was used as a measure of the protein’s importance in the BLAD interactome. In this study high scores in centrality measures were combined with autoimmune disease ontology terms, in order to find which proteins are important within the network and have associations with autoimmune diseases.

From the analysis of the BLAD interactome, the identification of 14 candidate BLAD-related proteins was achieved. The most intriguing fact is the literature verification of the actual implication of 50% of candidate BLAD-related proteins in these disorders. Hence, this piece of evidence suggests that the remaining proteins that have not been confirmed should be further investigated for their potential implication in BLADs. Remarkably, the proteins appear to be involved in several functions of the immune system and have associations with several autoimmune diseases. The findings of our literature survey revealed that among the 14 candidate proteins, 2 proteins **CASP8** (6) and **FADD** (64) are already known to be associated with autoimmune lymphoproliferative syndrome (ALPS) (6) (**Supplementary Figure 6, A**). Remarkably, **DAXX** is also likely involved in ALPS pathogenesis (65) (**Supplementary Figure 6, A**). Moreover, *Maia et.al*. (66) concluded that susceptibility to autoimmune thrombocytopenic purpura (ITP) relates with genes located very close to **MICA** or even MICA itself (**Supplementary Figure 6, B**). Furthermore, MICA has an abnormal expression on granulocytes of PNH patients (67) (**Supplementary Figure 6, C**).

Interestingly, **TRAF6** is implicated in ITP megakaryopoiesis (68) (**Supplementary Figure 6, B**) and **FYN** is overexpressed in T-cells of mice with ALPS (69) (**Supplementary Figure 6, A**). Moreover, in accordance with the study of *Guinn et al.* (1995) (70), high levels of **TP53** have been recorded in an ITP patient (**Supplementary Figure 6, B**). More recent evidence by *Yadav et al.* (71) in 2016, shows overexpression of TP53 in pernicious anemia (PA) patients – indicating TP53 as a primary apoptosis inducer in megaloblasts (**Supplementary Figure 6, D**).

The aforementioned verification of our findings suggests that the approach followed in this study is useful for the identification of candidate BLAD-related proteins. As shown in **Supplementary Figure 6**, 4 proteins are verified for ALPS (**Supplementary Figure 6, A**), 3 proteins for ITP (**Supplementary Figure 6, B**) and 1 protein for PNH (**Supplementary Figure 6, C**) and PA (**Supplementary Figure 6, D**), respectively.

The findings also highlight that through clustering analysis, 17 bridge proteins were identified as candidate proteins for BLADs’ relationships. Notably, the fact that 2 out of the 17 bridge proteins (EGFR and TRAF6) are common with those obtained through the BLAD interactome analysis, demonstrate that these two different approaches are complementary – having their own merit in identifying candidate BLAD-related proteins.

As stated in a previous study (72) and as demonstrated from our results, disease-related proteins are clustered together. It is worth mentioning that in the 1^st^ cluster there are 4 proteins associated with BLADs and 5 candidate BLAD-related proteins (Fig. 5, **cluster 1 – nodes with thick black border)**. These results point to the likelihood that these 5 candidate BLAD-related proteins may be implicated in those BLADs. Significantly, in the 3^rd^ cluster, **TRAF6** (Fig. 5, **cluster 3 – orange node with thick black border**) is a bridge protein of IL1RN and TNF, both associated with ITP, while the latter is associated with PNH as well (Fig. 5, **3^rd^ cluster**). Regarding the fact that ITP and PNH are associated with each other (15), it could be assumed that TRAF6 and the remaining bridge proteins (IL1R1, ST13 and ATP2A2) may play an essential role in the relationship of ITP and PNH pathogenesis.

In the 1^st^ cluster, a 3^rd^ order connection between SBDS and FASLG (**Supplementary Figure 4**) is worth investigating, due to the interesting results shown by their bridge proteins in combination with literature findings. *Calado et al.* (73) proposed that an SBDS mutation is associated with acquired aplastic anemia and leukocytes’ telomere shortening. In accordance with their study, the maintenance of telomere length depends on SBDS contribution – since leukocytes’ telomere shortening was observed in SBDS-deficient patients, as opposed to others with normal SBDS expression. Given the fact that the authors failed to find any physical associations among SBDS and telomerase complex components (TERT and TERC), they suggested that telomere length can be influenced by SBDS via a telomere-independent pathway. Nevertheless, as indicated from our results, SBDS interacts with TERF2IP (also known as RAP1) and POT1, which form, with 4 additional telomere-associated proteins (TPP1, TIN2, TRF1, and TRF2), the shelterin complex. Together, the shelterin complex and telomerase, are vital for telomeres’ maintenance (74). TERF2IP and TRF2 form a complex which affects the human telomeres’ length and heterogeneity – playing a role in telomere function (75). POT1 can act positively as a telomerase-dependent regulator of telomere length (76,77). Therefore, dysregulation of SBDS may affect telomere length, through TERF2IP and POT1.

A connection between severe aplastic anemia (SAA) and telomere length has been reported by *Wang et al.* (78), where low expression of *pot1* has been detected in SAA patients, whose telomeres were found to be short. Moreover, in these patients, proinflammatory cytokines, TNF-α and IFN-γ, are also found at high concentrations, suggesting that apoptosis or senescence triggering occurs via *pot1* and *atm/atr* activation. In light of recent evidence (79,80), inhibition of SBDS results in accelerated apoptosis through the Fas pathway. *Watanabe et al.* (80) proposed that SBDS’s role regarding ribosome biogenesis might be associated with proteins regulating apoptosis via the Fas pathway. Our findings point to the likelihood that SBDS may affect FASLG, an effector of apoptosis, through the proteins TERF2IP and POT1. This assumption seems to be possible, considering that *Lee et al.* (81) designated apoptosis as one of the top biological processes of the telomere interactome. Interestingly, they suggested that the aforementioned proteins of the shelterin complex may have a connecting role between telomere dysfunction and apoptosis. We hypothesize that the bridge proteins CRK, TRIP10 and PACSIN isoforms 1-3 (**Supplementary Figure 4**), could play a significant role in that process since they are adapter proteins with an essential role in signal transduction pathways. As it has been shown by other studies, CRK was identified as a pro-apoptotic protein (82), in addition to TRIP10 and PACSIN isoforms 1-3, known as the “FCH/SH3-family”, which were all found as interaction partners of the proline-rich region of FASLG – for safe regulation of its storage and transport (83).

In regard to the 2^nd^ order connection of PTPN22-SH2B3 (1^st^ cluster) (**Supplementary Figure 3**), the bridge proteins EGFR and ERBB2 were found to connect them. Our suggestion about possible ITP-APS association is in line with a previous study (16). Several studies have shown an association among APS, the premature formation of atherosclerotic plaques and the incident acceleration of atherosclerosis (84). Additionally, ITP patients present an increased risk for atherosclerosis (85). To the best of our knowledge, the transfer of peripheral monocytes to the subendothelial space, their differentiation into macrophages followed by proliferation, is important during atherosclerosis. It is worth noting that EGFR was found, among other cell types, on monocytes and macrophages present within the atherosclerotic plaque (86). Interestingly, epidermal growth factor (EGF), one of EGFR’s ligands, was identified at elevated levels in ITP patients with higher platelet counts, as compared to those with lower platelet counts (87).

HPO analysis highlighted common phenotypic abnormalities among bridge proteins and proteins associated with BLADs. This confirms previous findings in the literature (18) that highly interconnected proteins have similar phenotypes.

It is worth mentioning that diseases associated with proteins that interact with each other are more likely correlated. In particular, some autoimmune diseases associated with bridge proteins are found to be related to BLADs. Specifically, EGFR which interacts with SH2B3 associated with APS (**Supplementary Figure 2**), is related to systemic lupus erythematosus (SLE) (88). Surprisingly, an increasing number of studies (89–91) have found APS in patients with SLE. Also, SKAP2 bridge protein, which interacts with PTPN22 associated with ITP (**Supplementary Figure 2**), is related to Crohn disease (92). This was first reported by *Baudard et al.* (93) in a case documenting the comorbidity of ITP and Crohn disease. Remarkably, SKAP2 and PTPN22 (**Supplementary Figure 2**) are both associated with type I diabetes mellitus (94), and an association of ITP-type I diabetes mellitus was found (95).

In conclusion, this study provides a systemic perspective of the relationships of proteins associated with BLADs. Our approach combines information of network theory analysis and statistically significant autoimmune disease associations, allowing us to identify 14 candidate BLAD-related proteins, out of which 7 are verified through literature searches (**CASP8, FADD, DAXX, MICA, TRAF6, FYN**, and **TP53**). We propose that the remaining 7 proteins (**EGFR, CDC42, LRRK2, MET**, **FN1, KIT**, and **IKBKE)** are potentially involved in the pathogenesis of BLADs. Clustering analysis allowed us to find 17 bridge proteins which may play a “connector” role between proteins associated with BLADs, and possibly BLAD interconnections. From those, the most prominent are **EGFR**, **TRAF6**, **POT1**, **TERF2IP**, **CRK**, **TRIP10**, and **PACSIN** isoforms 1-3. It is worth noting that EGFR and TRAF6 have been detected as both candidate BLAD-related proteins and as bridge proteins. Our results are also confirmed from numerous experimental works. However, additional experimental studies, based on the results of our computational analysis, should be conducted, to investigate the roles of all these proteins in BLADs.

## Acknowledgement

The authors thank the National and Kapodistrian University of Athens for use of premises and equipment.

## Abbreviations

BLADs: Blood-cell targeting autoimmune diseases
PMIDs: PubMed IDs
PPI: Protein-protein interaction
MCL: Markov Clustering Algorithm
PHC: Proteins with high closeness centrality
HPO: Human Phenotype Ontology
CC: Clustering coefficient
CPL: Characteristic path length
ALPS: Autoimmune lymphoproliferative syndrome
PNH: Paroxysmal nocturnal hemoglobinuria
ITP: Autoimmune thrombocytopenic purpura
APS: Antiphospholipid syndrome
PA: Pernicious anemia
CASP8: Caspase-8
FADD: FAS-associated death domain protein
DAXX: Death domain-associated protein
MICA: MHC class I polypeptide-related sequence A
TRAF6: TNF receptor-associated factor 6
FYN: Tyrosine-protein kinase Fyn
TP53: Cellular tumor antigen p53
EGFR: Epidermal growth factor receptor
STAT5: Signal transducer and activator of transcription 5
HBEGF: Heparin-binding epidermal growth factor
SBDS: Ribosome maturation protein SBDS
FASLG: Tumor necrosis factor ligand superfamily member 6
PTPN22: Tyrosine-protein phosphatase non-receptor type 22
SH2B3: SH2B adapter protein 3
STAT5: signal transducer and activator of transcription 5
POT1: Protection of telomeres protein 1
TERF2IP: Telomeric repeat binding factor 2-interacting protein 1

