## Supplementary File 1 for "Visualization and Analysis of the Interaction Network of Proteins Associated with Blood-cell targeting Autoimmune Diseases"

**Web application User guide**

(<http://83.212.109.111/BLAD>)

In order to create an interactive network visualization, **Cytoscape.js** was used. Cytoscape.js, a graph theory library, allows for the display and manipulation of rich, interactive graphs such as the BLAD interactome. For more details please refer to Fig 2, 4 and 5 of the main manuscript. In the main window the user can see the entire network and “interact” with it.

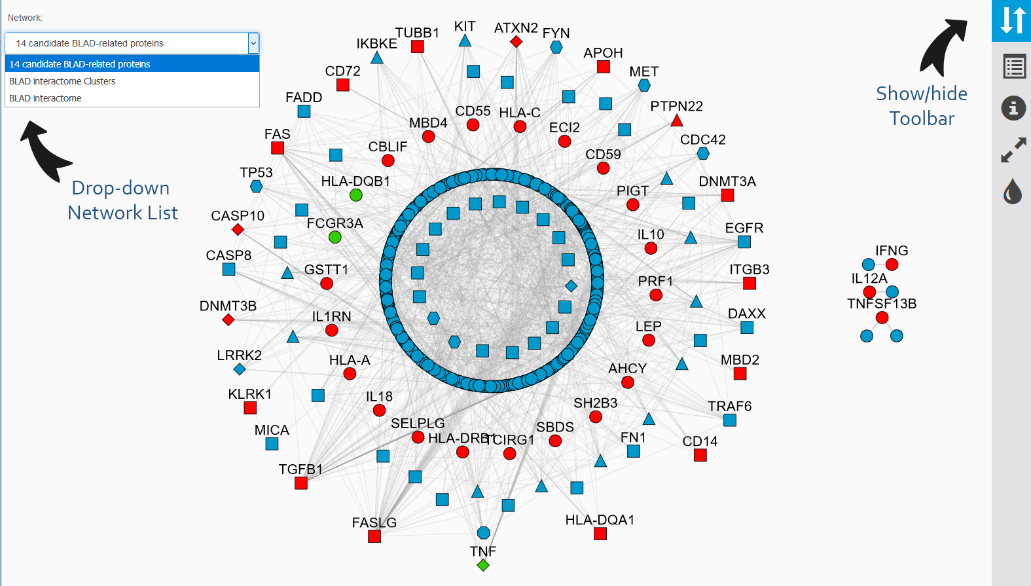

The user is allowed to select between the different networks described in the main manuscript: ***Fig. 2*** (**The BLAD Interactome**), ***Fig. 4*** (**14 candidate BLAD-related proteins**) and the clustered network of ***Fig. 5*** (**BLAD interactome clusters**), using the drop-down network list on the upper left corner of the window as shown above.

A button on the upper right corner of the screen allows the user to **show/hide the toolbar** of

|  | 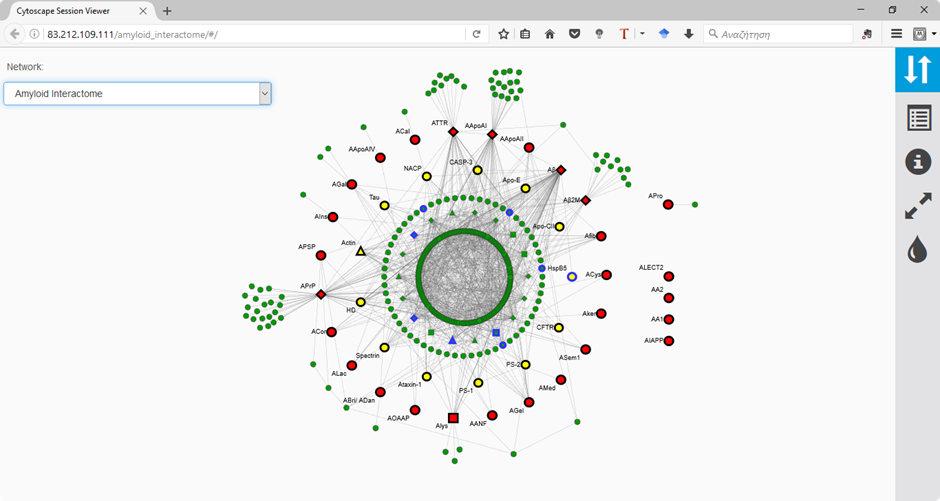 | the application. Among the available functions, the **fit** to window button lets the user | | | | | |
| --- | --- | --- | --- | --- | --- | --- | --- |
|  | to adjust the network to its original position | | 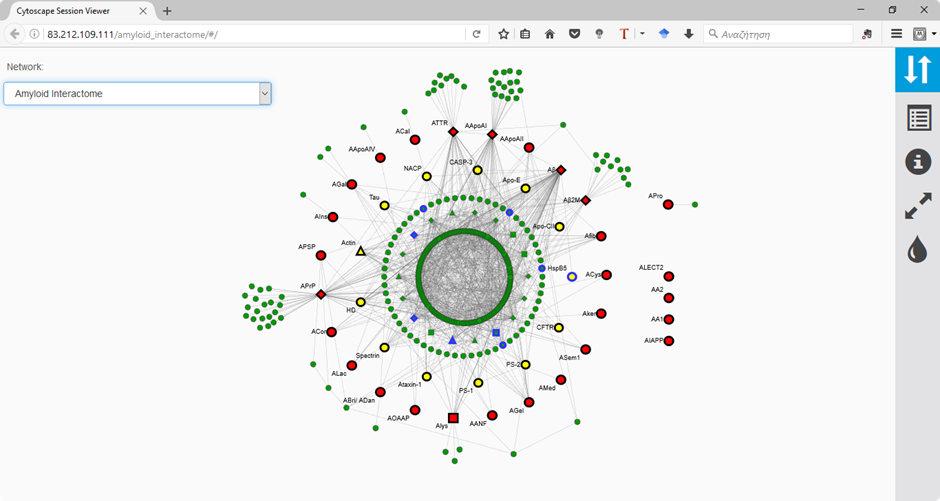 | whereas the background colour allows the | | | |
|  | change of the network’s background appearance. | | | 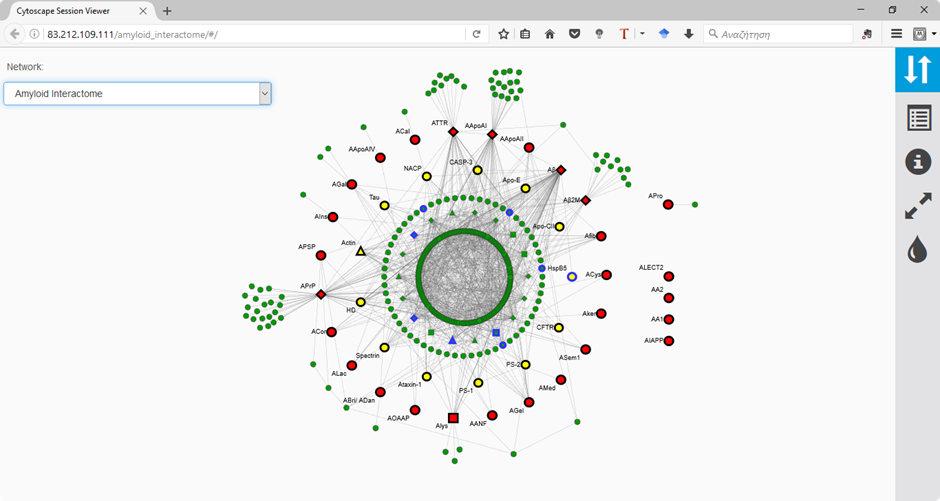 |  | | |
| Important information can be gathered from the Table Pane | | | | | | 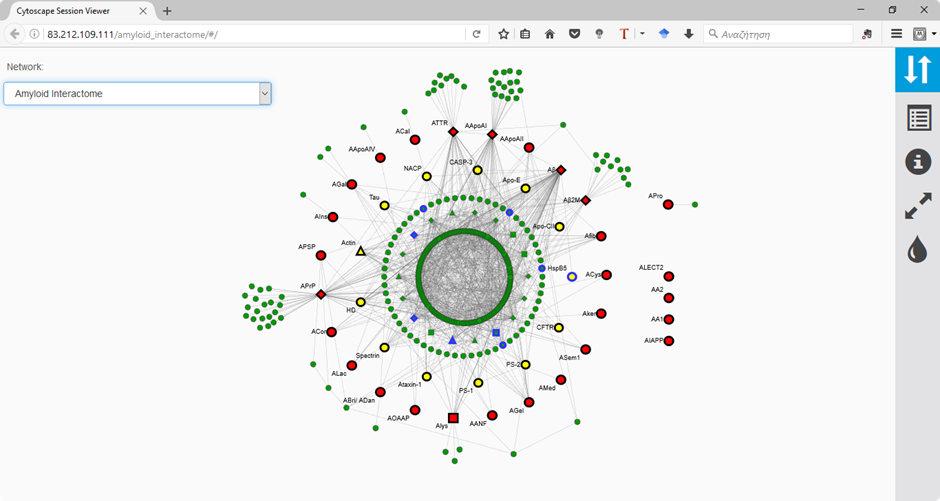 | where the user can see |
| all relevant information regarding selected nodes. | | | | | | | |

Specifically, the **Node Table** contains **unique identifiers** for each node (UniProt AC), **Protein Names, Gene Names** and **Function** for all proteins. It also contains all **Network Centrality** measures (e.g. node degree centrality, betweenness centrality etc.) for each protein as described in the paper. For **proteins of interest** a column with **the preferable Gene Name**, used in the main manuscript, is given. Part of the Node Table is shown below.

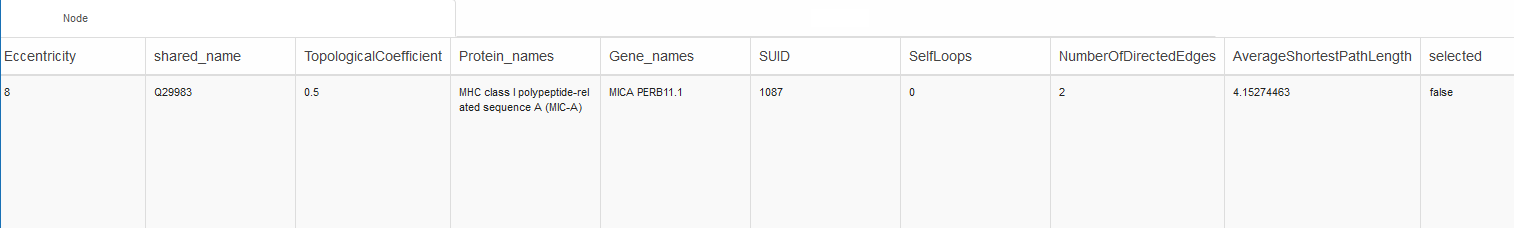
