## Supplementary material for "Visualization and Analysis of the Interaction Network of Proteins Associated with Blood-cell targeting Autoimmune Diseases": Table 1

**Table 1:** List of Blood-cell targeting Autoimmune Diseases (BLADs) with the corresponding target cells, description/symptoms, alternative names and PMIDs.

| **BLADs** | **Target** | **Description/Symptoms** | **Alternative names** | **PMIDs** |
| --- | --- | --- | --- | --- |
| Antiphospholipid syndrome | platelets | Presence of antibodies directed against phospholipids. Condition associated with a variety of diseases, notably systemic lupus erythematosus and other connective tissue diseases, thrombopenia, and arterial or venous thromboses. | Antiphospholipid antibody syndrome (APS); APLS; Anticardiolipin syndrome; Anticardiolipin antibody syndrome; aCL syndrome; aPL syndrome; Hughes syndrome; Lupus anticoagulant syndrome; Anti-Phospholipid Antibody Syndrome; Anti-Phospholipid Syndrome | 27024977;26858847; 16972844;9814662; 22635209;8712801 |
| Acquired aplastic anemia | hematopoietic cells | Immune-mediated reduction of hematopoiesis. Dysregulation of T-cell homeostasis and abnormal production of cytokines including tumor necrosis factor-α, interferon-γ and transforming growth factor-β induce apoptosis of hematopoietic stem/progenitor cells. | - | - |
| Autoimmune hemolytic anemia | erythrocytes | Acquired hemolytic anemia due to the presence of autoantibodies which agglutinate or lyse red blood cells. | Autoimmune hemolytic anemia (AIHA); Autoimmune haemolytic disease (cold type) (warm type); Acquired Autoimmune Hemolytic Anemia; Cold Agglutinin Disease; Cold Antibody Disease; cold antibody autoimmune hemolytic anemia; Cold Antibody Hemolytic Anemia; Idiopathic Autoimmune Hemolytic Anemia; Paroxysmal Cold Hemoglobinuria; Idiopathic Acquired Hemolytic Anemia | 24385063;6034957; 19074065;27879544 |
| Autoimmune lymphoproliferative syndrome | lymphocytes | Rare congenital lymphoid disorder due to mutations in certain Fas-Fas ligand pathway genes. Known causes include mutations in FAS, TNFSF6, NRAS, CASP8, and CASP10 proteins. Clinical features include lymphadenopathy, splenomegaly and autoimmunity. | Autoimmune Lymphoproliferative Syndrome Type 1-Autosomal Dominant; Autoimmune Lymphoproliferative Syndrome Type I- Autosomal Dominant; Autoimmune Lymphoproliferative Syndrome Type 2B; Autoimmune Lymphoproliferative Syndrome Type 2B (ALPS2B); Autoimmune Lymphoproliferative Syndrome Type Iib; Canale Smith Syndrome; Canale-Smith Syndrome | 17999750;14749982 |
| Autoimmune neutropenia | neutrophils | Autoantibodies directed against membrane antigens of neutrophils cause their increased peripheral destruction. | Autoimmune neutropaenia; Immunologic neutropenia; Acquired neutropenia | 15561677 |
| Autoimmune thrombocytopenic purpura | platelets | Thrombocytopenia occurring in the absence of toxic exposure or any disease associated with decreased platelet counts. In most cases mediated by immunoglobulin G autoantibodies, which attach to platelets and lead to their destruction by macrophages. Disease presented in acute (pediatric) and chronic (adult) forms. | Evans syndrome; Evans' Syndrome; Autoimmune Thrombocytopenia; Immune Thrombocytopenia; Immune Thrombocytopenic Purpura; ITP; Werlhof Disease; Werlhof's Disease; primary thrombocytopenic purpura; primary immune thrombocytopenic purpura; primary immune thrombocytopenia; idiopathic thrombocytopenic purpura | 23273499;27861740; 19444418;27429531; 9167472 |
| Pernicious anemia | erythrocytes | Megaloblastic anemia occurring in children, but more commonly in adults. Characterized by histamine-fast achlorhydria. Laboratory and clinical manifestations are based on malabsorption of vitamin B 12 due to failure of the gastric mucosa to secrete adequate and potent intrinsic factor. | Addison's Anemia; Addison-Biermer Anemia; Addisonian Pernicious Anemia | 25787024 |
| Primary acquired pure red cell aplasia | erythroids | Interruption of erythroid differentiation usually mediated by an autoantibody. Characterized by marked reduction or absence of red blood cell precursors (reticulocytes) from the bone marrow. | Primary autoimmune PRCA (includes transient erythroblastopenia of childhood); Primary myelodysplastic PRCA | 27913462 |
| Paroxysmal nocturnal hemoglobinuria | erythrocytes | Characterized by recurrence of hemoglobinuria caused by intravascular hemolysis. In cases occurring during or after sleep (paroxysmal nocturnal hemoglobinuria), clonal hematopoietic stem cells exhibit global deficiency of cell membrane proteins. | PNH; Marchiafava-Micheli syndrome; Marchiafava-Micheli Syndrome | 2097747;17285400 |
