## Supplementary material for "Visualization and Analysis of the Interaction Network of Proteins Associated with Blood-cell targeting Autoimmune Diseases": Table 2

**Table 2**: Data set of proteins associated with BLADs.

| **BLAD** | **UniProt AC** | **Gene** | **Protein name** |
| --- | --- | --- | --- |
| Antiphospholipid syndrome | P02749 | APOH | Beta-2-glycoprotein 1 |
|  | Q14242 | SELPLG | P-selectin glycoprotein ligand 1 |
|  | Q99700 | ATXN2 | Ataxin-2 |
|  | Q9UQQ2 | SH2B3 | SH2B adapter protein 3 |
| Acquired aplastic anemia | P30711 | GSTT1 | Glutathione S-transferase theta-1 |
|  | P09488 | GSTM1 | Glutathione S-transferase Mu 1 |
|  | P14222 | PRF1 | Perforin-1 |
|  | O75521 | ECI2 | Enoyl-CoA delta isomerase 2, mitochondrial |
|  | P07099 | EPHX1 | Epoxide hydrolase 1 |
|  | Q9Y3A5 | SBDS | Ribosome maturation protein SBDS |
|  | Q8TDQ0 | HAVCR2 | Hepatitis A virus cellular receptor 2 |
| Autoimmune hemolytic anemia | - | - | - |
| Autoimmune lymphoproliferative syndrome | P25445 | FAS | Tumor necrosis factor receptor superfamily member 6 |
|  | P48023 | FASLG | Tumor necrosis factor ligand superfamily member 6 |
|  | Q92851 | CASP10 | Caspase-10 |
| Autoimmune neutropenia | P01920 | HLA-DQB1 | MHC class II antigen DQB1 |
| Autoimmune thrombocytopenic purpura | P60568 | IL2 | Interleukin-2 |
|  | P18510 | IL1RN | Interleukin-1 receptor antagonist protein |
|  | Q14116 | IL18 | Interleukin-18 |
|  | O95998 | IL18BP | Interleukin-18-binding protein |
|  | P29459 | IL12A | Interleukin-12 subunit alpha |
|  | Q9Y275 | TNFSF13B | Tumor necrosis factor ligand superfamily member 13B |
|  | P21854 | CD72 | B-cell differentiation antigen CD72 |
|  | Q9Y6K1 | DNMT3A | DNA (cytosine-5)-methyltransferase 3A |
|  | Q9UBC3 | DNMT3B | DNA (cytosine-5)-methyltransferase 3B |
|  | P23526 | AHCY | Adenosylhomocysteinase |
|  | Q8NEV9 | IL27 | Interleukin-27 subunit alpha |
|  | Q9Y2R2 | PTPN22 | Tyrosine-protein phosphatase non-receptor type 22 |
|  | Q9UBB5 | MBD2 | Methyl-CpG-binding domain protein 2 |
|  | O95243 | MBD4 | Methyl-CpG-binding domain protein 4 |
|  | P41159 | LEP | Leptin |
|  | Q13488 | TCIRG1 | V-type proton ATPase 116 kDa subunit a isoform 3 |
|  | P22301 | IL10 | Interleukin-10 |
|  | Q9H4B7 | TUBB1 | Tubulin beta-1 chain |
|  | P08637 | FCGR3A | Low affinity immunoglobulin gamma Fc region receptor III-A |
|  | P01375 | TNF | Tumor necrosis factor (TNF-alpha) |
|  | Q8TDQ0 | HAVCR2 | Hepatitis A virus cellular receptor 2 |
| Autoimmune thrombocytopenic purpura in systemic lupus erythematosus | P05106 | ITGB3 | Integrin beta-3 |
| Pernicious anemia | P27352 | GIF | Gastric intrinsic factor |
| Primary acquired pure red cell aplasia | - | - | - |
| Paroxysmal nocturnal hemoglobinuria | P37287 | PIGA | Phosphatidylinositol N-acetylglucosaminyltransferase subunit A |
|  | P01911 | HLA-DRB1 | HLA class II histocompatibility antigen, DRB1-15 beta chain |
|  | P01909 | HLA-DQA1 | MHC class II DQA1 |
|  | P01892 | HLA-A | HLA class I histocompatibility antigen, A-2 alpha chain |
|  | P30462 | HLA-B | HLA class I histocompatibility antigen, B-14 alpha chain |
|  | P30505 | HLA-C | HLA class I histocompatibility antigen, Cw-8 alpha chain |
|  | P04229 | HLA-DRB1 | HLA class II histocompatibility antigen, DRB1-1 beta chain |
|  | Q969N2 | PIGT | GPI transamidase component PIG-T |
|  | P01137 | TGFB1 | Transforming growth factor beta-1 |
|  | P01579 | IFNG | Interferon gamma |
|  | P26718 | KLRK1 | NKG2-D type II integral membrane protein |
|  | P13987 | CD59 | CD59 glycoprotein |
|  | P08174 | CD55 | Complement decay-accelerating factor |
|  | P08571 | CD14 | Monocyte differentiation antigen CD14 |
|  | P01920 | HLA-DQB1 | MHC class II antigen DQB1 |
|  | P08637 | FCGR3A | Low affinity immunoglobulin gamma Fc region receptor III-A |
|  | P01375 | TNF | Tumor necrosis factor (TNF-alpha) |
