## Supplementary material for "Visualization and Analysis of the Interaction Network of Proteins Associated with Blood-cell targeting Autoimmune Diseases": Table 3

**Table 3:** **Identification of candidate BLAD-related proteins**. The column “**Centrality measures”** presents proteins which are ranked in various centrality measures in the order: *Degree-D, Betweenness-B* and *Closeness-C*. The rank of each protein is given inside the braces of the corresponding centrality measure in the top 48 rankings (approximately the top 10% of the network's proteins). Proteins ranking high in all three measures are shown first, followed by proteins with two measures and one measure, respectively. The column **“Autoimmune disease ontology terms”** represents the statistically significant autoimmune disease ontology terms per protein. The column “**Interactions with proteins associated with BLADs”** shows the interaction(s) of each protein with protein(s) associated with BLADs - whereas the corresponding association with each BLAD can be seen inside the braces.

| **UniProt AC** | **Gene** | **Protein name** | **Centrality**  **measures**  **(Ranking)** | **Autoimmune disease ontology terms** | **Interactions with proteins associated with BLADs** |
| --- | --- | --- | --- | --- | --- |
| P00533 | *EGFR* | Epidermal growth factor receptor | D (3), B (4), C (9) | Systemic lupus erythematosus | PTPN22 (autoimmune thrombocytopenic purpura),  TGFB1 (paroxysmal nocturnal hemoglobinuria),  CD59 (paroxysmal nocturnal hemoglobinuria),  SH2B3 (antiphospholipid syndrome) |
| Q14790 | *CASP8* | Caspase-8 | D (17), B (20), C (26) | Multiple sclerosis | FAS (autoimmune lymphoproliferative syndrome),  FASLG (autoimmune lymphoproliferative syndrome),  CASP10 (autoimmune lymphoproliferative syndrome) |
| Q9Y4K3 | *TRAF6* | TNF receptor-associated factor 6 | D (21), B (19), C (31) | Rheumatoid arthritis, Crohn's disease, Ulcerative colitis | AHCY (autoimmune thrombocytopenic purpura) |
| Q9UER7 | *DAXX* | Death domain-associated protein 6 | D (35), B (29), C (27) | Systemic lupus erythematosus, Type I diabetes mellitus | TGFB1 (paroxysmal nocturnal hemoglobinuria),  FAS (autoimmune lymphoproliferative syndrome) |
| P60953 | *CDC42* | Cell division control protein 42 homolog | D (36), C (40) | Relapsing-Remitting Multiple sclerosis | APOH (antiphospholipid syndrome) |
| Q5S007 | *LRRK2* | Leucine-rich repeat serine/threonine-protein kinase 2 | D (44), B (41) | Crohn's disease | TUBB1 (autoimmune thrombocytopenic purpura) |
| P04637 | *TP53* | Cellular tumor antigen p53 | D (46), C (45) | Systemic lupus erythematosus, Multiple sclerosis, Type I diabetes mellitus | APOH (antiphospholipid syndrome) |
| P06241 | *FYN* | Tyrosine-protein kinase Fyn | D (28), C (17) | Systemic lupus erythematosus | FAS (autoimmune lymphoproliferative syndrome),  FASLG (autoimmune lymphoproliferative syndrome) |
| P08581 | *MET* | Hepatocyte growth factor receptor | D (20), C (33) | Multiple sclerosis | SH2B3 (antiphospholipid syndrome) |
| Q29983 | *MICA* | MHC class I polypeptide-related sequence A | B (33) | Systemic lupus erythematosus, Multiple sclerosis, Celiac disease, Psoriasis, Graves disease, Behcet syndrome, Addison disease, Type I diabetes mellitus | KLRK1 (paroxysmal nocturnal hemoglobinuria) |
| Q13158 | *FADD* | FAS-associated death domain protein | C (37) | Systemic lupus erythematosus, Type I diabetes mellitus | FAS (autoimmune lymphoproliferative syndrome),  FASLG (autoimmune lymphoproliferative syndrome),  CASP10 (autoimmune lymphoproliferative syndrome),  MBD4 (autoimmune thrombocytopenic purpura) |
| P02751 | *FN1* | Fibronectin | C (38) | Systemic sclerosis, Type I diabetes mellitus | FASLG (autoimmune lymphoproliferative syndrome) |
| P10721 | *KIT* | Mast/stem cell growth factor receptor Kit | D (39) | Vitiligo | SH2B3 (antiphospholipid syndrome) |
| Q14164 | *IKBKE* | Inhibitor of nuclear factor kappa-B kinase subunit epsilon | D (45) | Rheumatoid arthritis | SBDS (acquired aplastic anemia) |
