## Supplementary File 2 for "Visualization and Analysis of the Interaction Network of Proteins Associated with Blood-cell targeting Autoimmune Diseases"

Assistant Prof. Vassiliki A. Iconomidou

Section of Cell Biology and Biophysics, Department of Biology, School of Sciences, National and Kapodistrian University of Athens, Athens 15701.

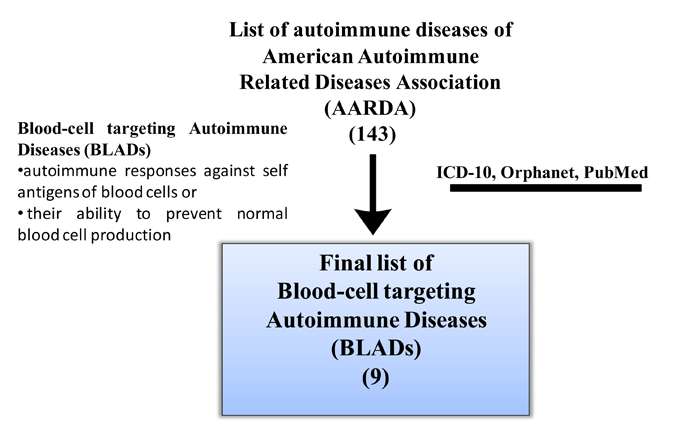

**Supplementary Figure 1. An overview of the basic protocol, used to collect blood-cell targeting autoimmune diseases (BLADs).**

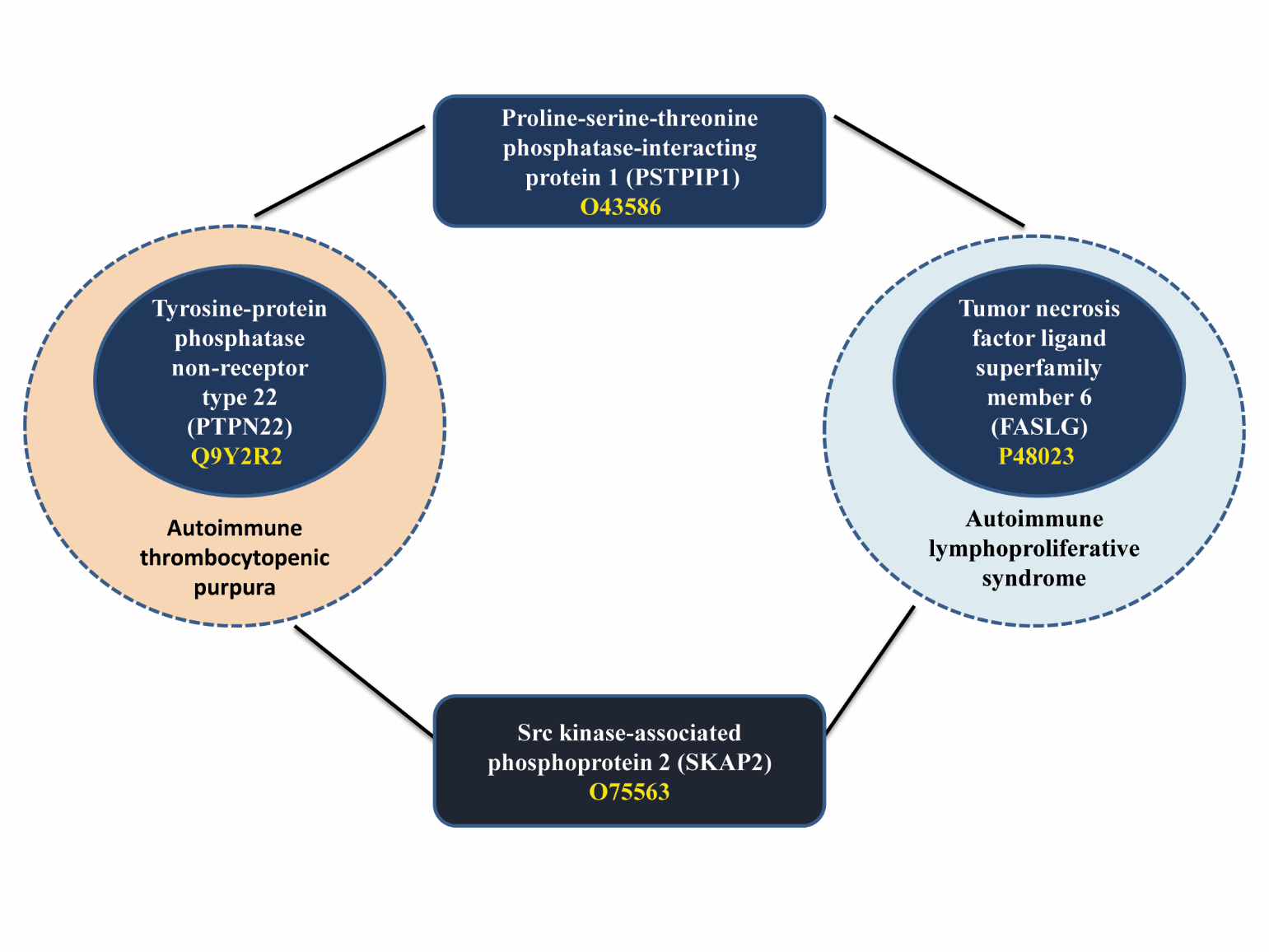

**Supplementary Figure 2: The 2^nd^ order connection among PTPN22-FASLG (1^st^ cluster).** The bridge proteins are PSTPIP1 and SKAP2. For each protein, the protein name, the gene name inside the braces, and the UniProt AC are given.

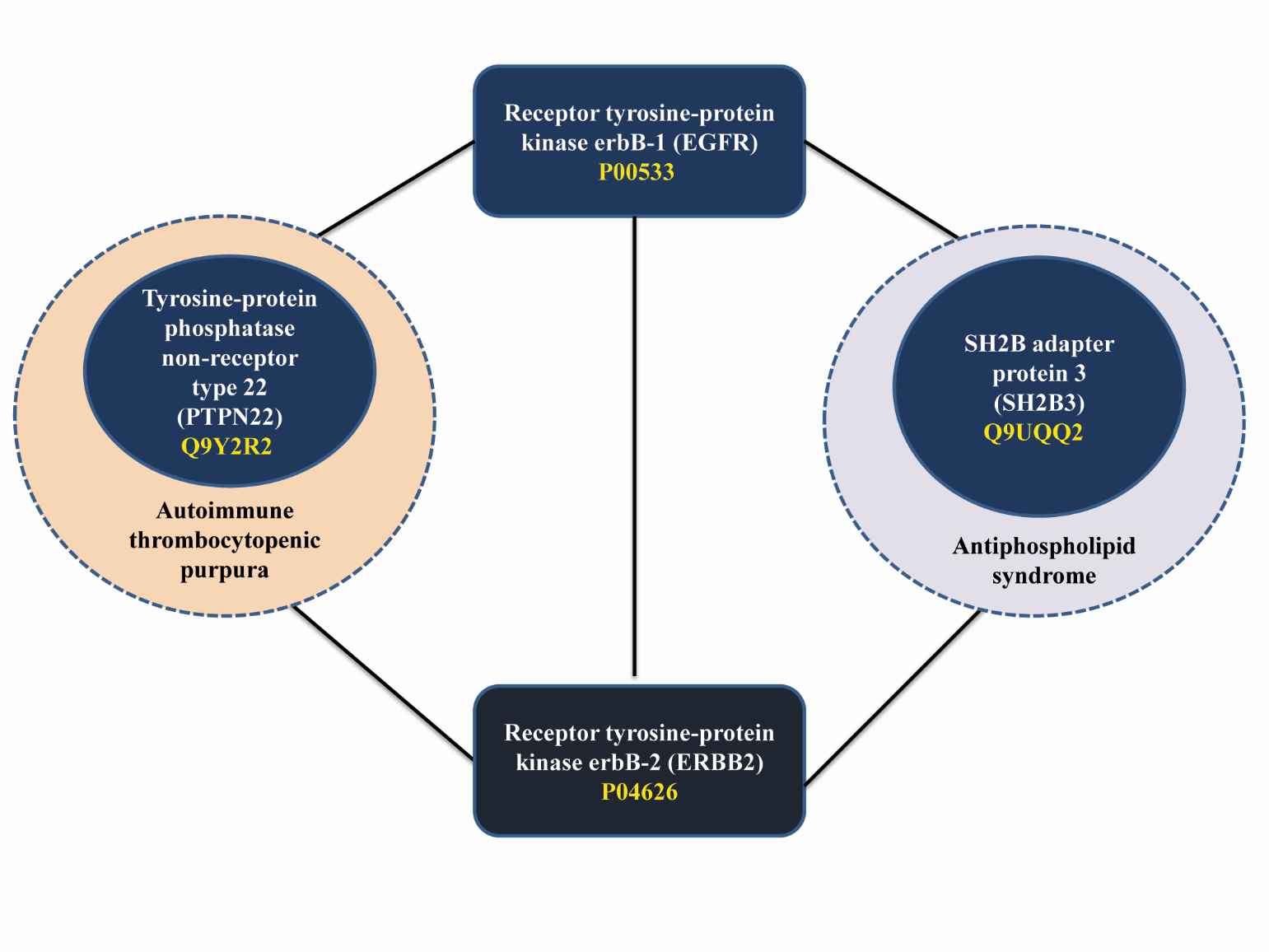

**Supplementary Figure 3: The 2^nd^ order connection among PTPN22-SH2B3 (1^st^ cluster).** The bridge proteins are EGFR and ERBB2. For each protein, the protein name, the gene name inside the braces, and the UniProt AC are given.

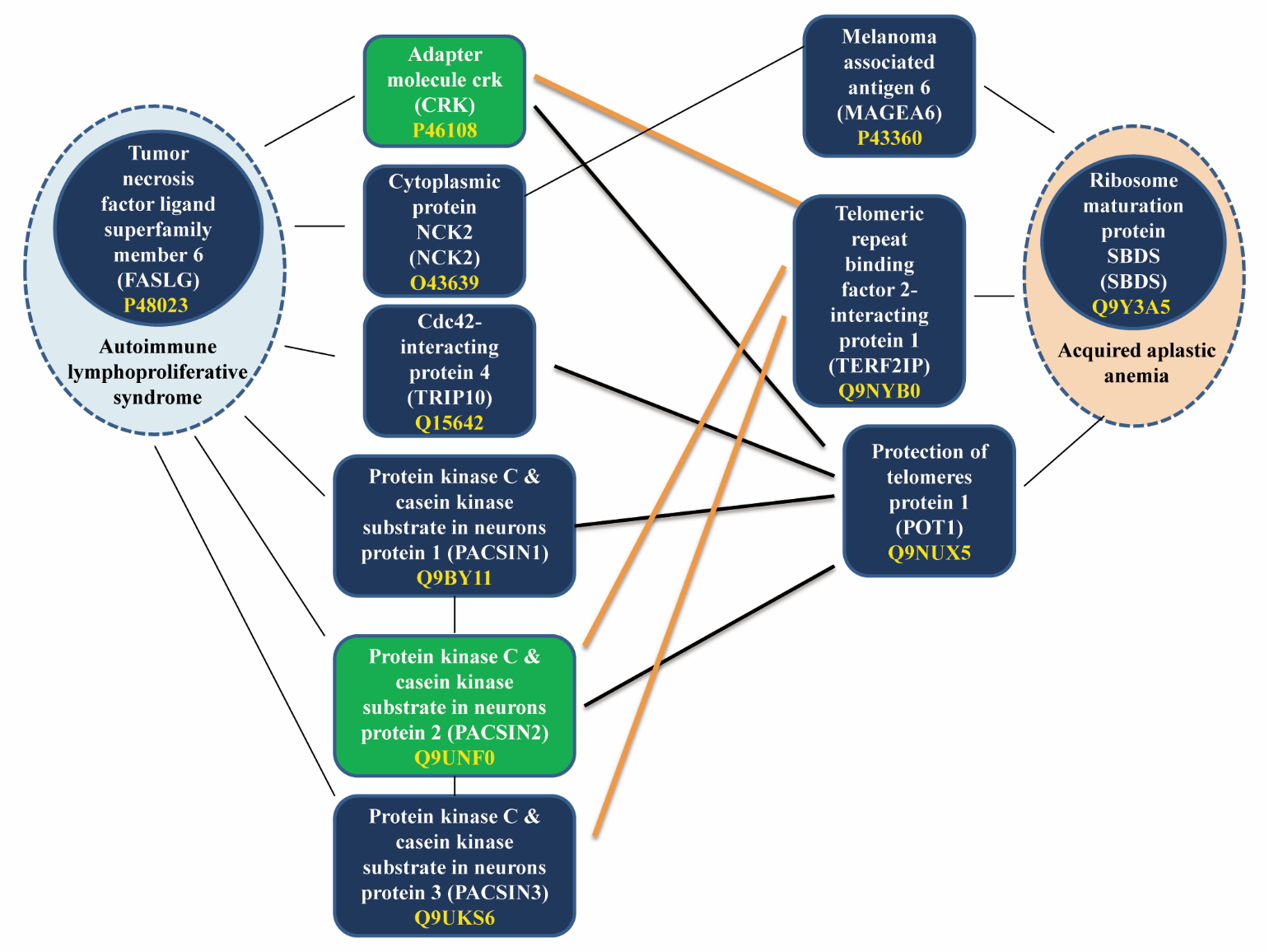

**Supplementary Figure 4: The 3^rd^ order connection among FASLG-SBDS (1^st^ cluster).** The bridge proteins are CRK, NCK2, TRIP10, PACSIN1, PACSIN2, PANCSIN3, MAGEA6, TERF2IP and POT1. Green-colored nodes represent proteins which interact with TERF2IP and POT1. The thick black lines depict the interaction between POT1 and other proteins. The thick orange lines depict the interaction between TERF2IP and other proteins. For each protein, the protein name, the gene name inside the braces, and the UniProt AC are given.

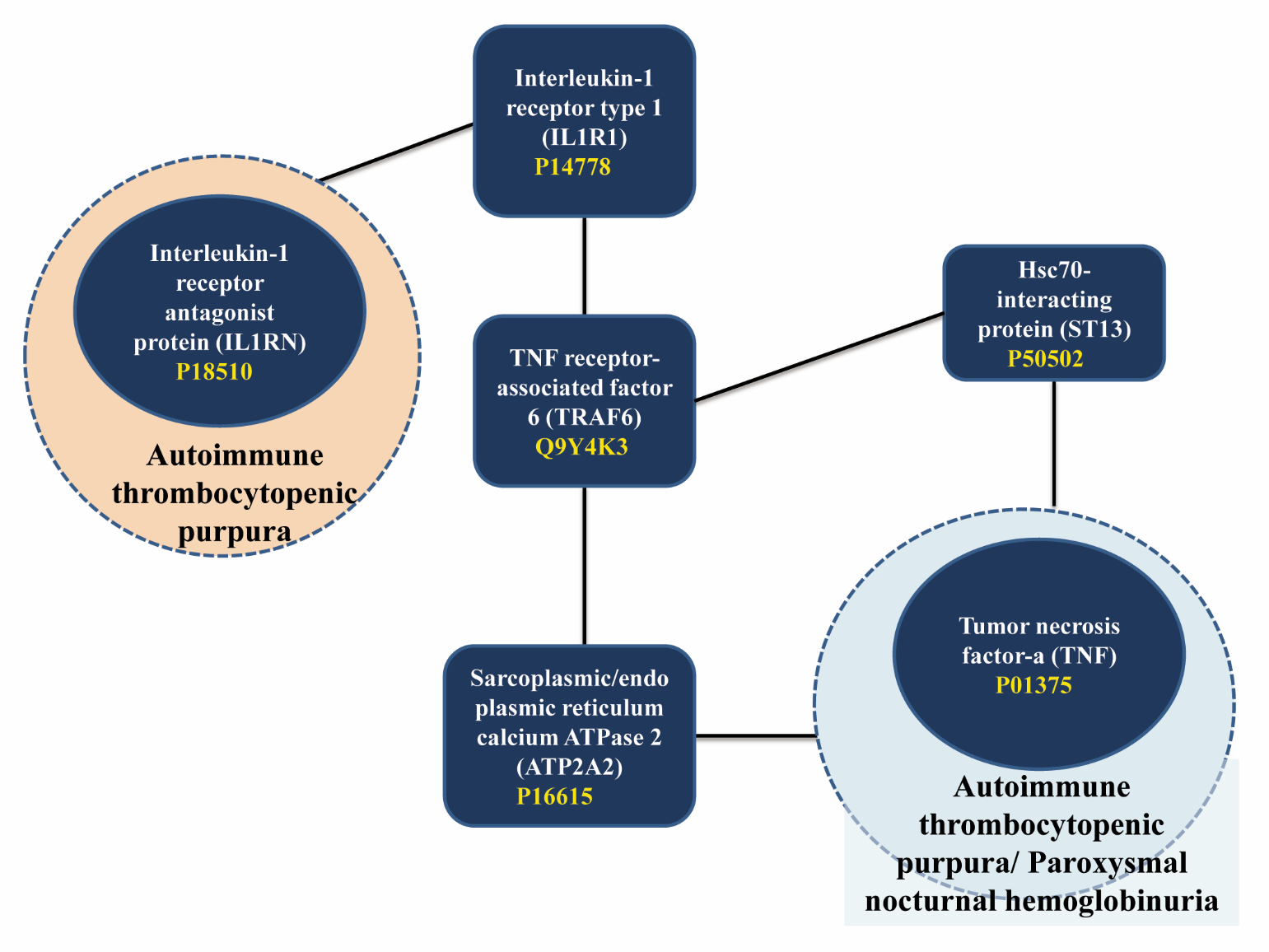

**Supplementary Figure 5: The 4^th^ order connection among IL1RN-TNF (3^rd^ cluster**). The bridge proteins are IL1R1, TRAF6, ATP2A2 and ST13. For each protein, the protein name, the gene name inside the braces, and the UniProt AC are given.

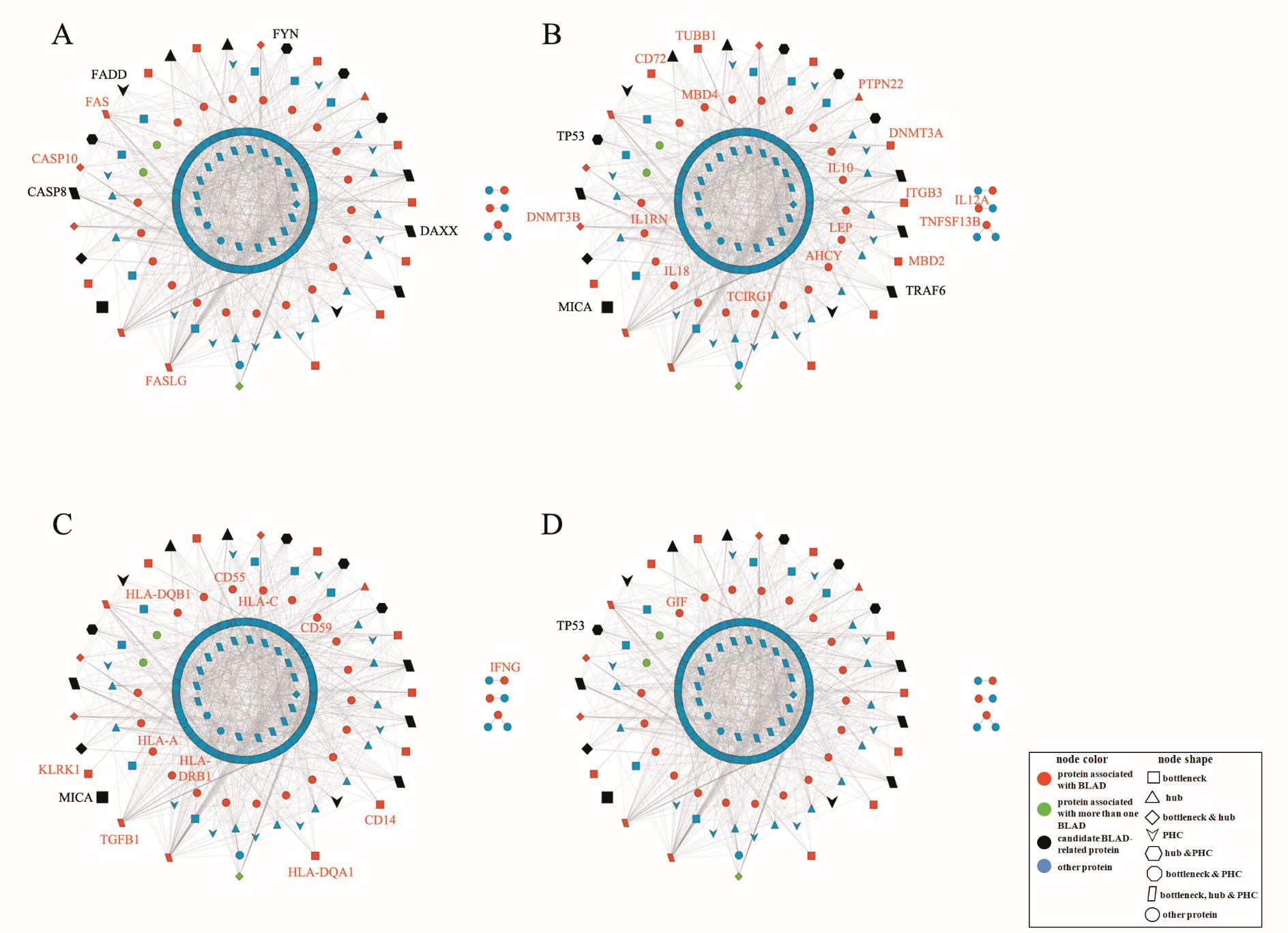

**Supplementary Figure 6: Proteins associated with (A) autoimmune lymphoproliferative syndrome (ALPS), (B) autoimmune thrombocytopenic purpura (ITP), (C) paroxysmal nocturnal hemoglobinuria (PNH) and (D) pernicious anemia (PA).** Proteins marked in red are the ones that belong to our initial data set (proteins associated with BLADs). Proteins marked in black are those identified as candidate BLAD-related proteins and are literature-verified for their association with the aforementioned autoimmune diseases.
